# Programmatic access to bacterial regulatory networks with *regutools*

**DOI:** 10.1101/2020.04.29.068551

**Authors:** Joselyn Chávez, Carmina Barberena-Jonas, Jesus E. Sotelo-Fonseca, José Alquicira-Hernández, Heladia Salgado, Leonardo Collado-Torres, Alejandro Reyes

## Abstract

**Summary:** *RegulonDB* has collected, harmonized and centralized data from hundreds of experiments for nearly two decades and is considered a point of reference for transcriptional regulation in *Escherichia coli* K12. Here, we present the *regutools R* package to facilitate programmatic access to *RegulonDB* data in computational biology. *regutools* gives researchers the possibility of writing reproducible workflows with automated queries to *RegulonDB*. The *regutools* package serves as a bridge between *RegulonDB* data and the *Bioconductor* ecosystem by reusing the data structures and statistical methods powered by other *Bioconductor* packages. We demonstrate the integration of *regutools* with *Bioconductor* by analyzing transcription factor DNA binding sites and transcriptional regulatory networks from *RegulonDB*. We anticipate that *regutools* will serve as a useful building block in our progress to further our understanding of gene regulatory networks.

**Availability and Implementation:** *regutools* is an *R* package available through *Bioconductor* at bioconductor.org/packages/regutools.

**Contact:** github.com/ComunidadBioInfo/regutools.

## 1 Introduction

Bacteria are able to sense transient environmental changes by, for example, detecting extracellular metabolites (Seshasayee *et al*., 2006). To maintain homeostasis in such changing environments, cells switch the state of transcriptional regulators from active to inactive, or viceversa. These switches result in expression changes of the regulators’ gene targets (Ledezma-Tejeida *et al*., 2019). A classical example of a transcriptional response caused by an environmental stimuli is the *lac* operon regulatory circuit. In this circuit, the expression of three genes is induced when the cell senses a high concentration of lactose in the extracellular environment. The functional activity of these genes results in the import of lactose into the cytoplasm and the cleavage of lactose into glucose and galactose. The circuit is completed when the cell senses low concentrations of lactose and the expression of the three genes is turned off.

To understand each of these regulatory networks and their interactions as a biological system, the database *RegulonDB* has integrated, curated and harmonized data from classic molecular biology and high-throughput experiments to create the most comprehensive regulatory map to date of *Escherichia coli K12* (Santos-Zavaleta *et al*., 2018). Thanks to *RegulonDB*, researchers have been able to access and analyze these data altogether and even define new concepts of transcriptional regulation, such as the genetic sensory response unit (Ledezma-Tejeida *et al*., 2017). To date, *RegulonDB* is a highly used database with more than 1,200 citations (Gama-Castro *et al*., 2015).

To facilitate access to *RegulonDB* from a data analysis programming language, we present an *R/Bioconductor* package called *regutools*. The package *regutools* enables users to query *RegulonDB* and download data with a few lines of code. *regutools* also provides functions with the most popular queries to the database. The output of the queries are data structures of the *Bioconductor* ecosystem where the metadata of the database, such as the database versions, are also stored (Fig. 1). We anticipate that *regutools* will facilitate the integration of data from *RegulonDB* into the analysis of high-throughput experiments while enhancing the reproducibility of such analyses.

**Fig. 1.**
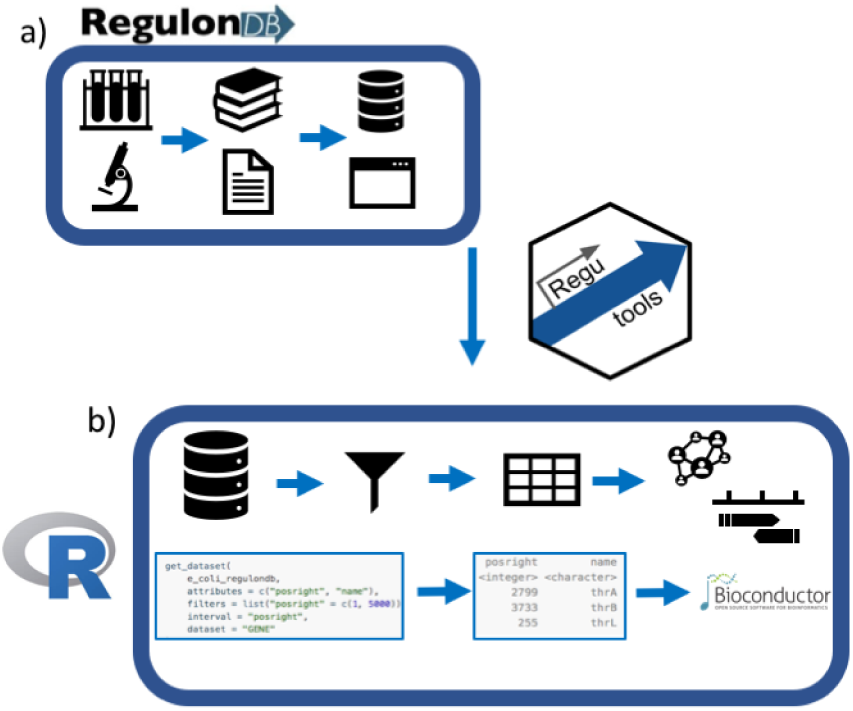
regutools overview: a) gene regulation experiments are curated into the RegulonDB database and are available through a web interface. b) regutools provides an interface to analyze the data with R and Bioconductor.

## 2 Implementation

The general paradigm of the *regutools* package is organized in three groups of functions that perform the following operations: (1) establish a connection to the database, (2) query the database and (3) integrate the results of the queries with other *R/Bioconductor* packages.

### Establishing a connection to the database

The original implementation of *RegulonDB* is a normalized relational *SQL* database. To ease integration with *R* and *Bioconductor*, we prepared a light-weight version of the database in *SQLite* format and made it available through *AnnotationHub*. In the *R* package, we designed an *S4* class called *regulondb*, which contains a connection to the database as well as the corresponding metadata such as organism name and versions of both the database and the reference genome. We provide a constructor function of the *regulondb* class that validates structure of the *SQLite* database and opens the connection to it. Note that the *regutools* implementation allows users to explore different versions of the *RegulonDB* database, including versions of other organisms that may be released in the future by *RegulonDB*.

### Querying the database

The general framework to query a *regulondb* object mimics the grammar of *Bioconductor* packages to access databases, such as *biomaRt* (Durinck *et al*., 2005). Specifically, a query consists of three elements: (1) a dataset of the database, (2) the attributes to retrieve and (3) a list of conditions to filter the results. For example, a user can query a table of the database (i.e. the dataset) to retrieve the gene names, genomic coordinates, and promoter sequences (i.e. the attributes) that are regulated by the Lacl repressor (i.e. the filter). In order to facilitate the process of designing a query, we implemented functions that retrieve the names and descriptions of all possible attributes and datasets. Additionally, the *regutools* package enables users to design complex filters, such as filtering by numeric ranges, partial matching, and using logical operators and multiple conditions.

The results of *regutools* queries are stored as an *S4* class called *regulondb_results* that is designed to also store the database metadata. The database metadata is automatically populated when the results of a query are generated.

Furthermore, the *regutools* package provides implementation of functions for the most popular queries to *RegulonDB*. These functions include retrieving: the names of the gene targets of a transcription factor (TF), the effect exerted in gene expression (i.e. whether the TF activates or represses expression, or both depending on the context), all DNA-binding sites of a TF, and all DNA elements that overlap between specified genomic coordinates.

### Integration with the *Bioconductor* ecosystem

To facilitate orchestration with *Bioconductor*, we implemented functions to convert *regulondb_results* objects to *Bioconductor*’s core objects. For example, a *regulondb_results* object can be converted to *GRanges* and *Biostrings* objects (Lawrence *et al*., 2013) whenever the results are genomic coordinates or sequences, respectively. Furthermore, we exemplify in the documentation how *regutools* queries can be integrated with other data analysis methods available in *Bioconductor*. For example, we demonstrate how DNA elements can be visualized in genome graphs using the package *Gviz* and how regulatory networks can be visualized using the *R* package *RCy3* (Hahne and Ivanek, 2016; Gustavsen *et al*., 2019).

## 3 Discussion

In the analysis of high-throughput experiments, integration from different sources of data is crucial for transforming data into biological knowledge. *regutools* provides a bridge between data from thousands of experiments harmonized by *RegulonDB* to *Bioconductor*, which provides a comprehensive ecosystem of data structures and statistical methods to analyse high-throughput experiments (Huber *et al*., 2015). We foresee that *regutools* will enable the scientific community to integrate, combine and re-analyze *RegulonDB* data using reproducible and reusable code.

## Acknowledgements

We thank Luis J. Muñiz-Rascado for releasing the *RegulonDB* database in *SQLite* format, Víctor Del Moral and César Bonavides for computational support in the lab, and Julio Collado-Vides for his advice and financial support.

## References

Durinck, S. et al. (2005). Biomart and bioconductor: a powerful link between biological databases and microarray data analysis. Bioinformatics, 21(16), 3439–3440.

Gama-Castro, S. et al. (2015). RegulonDB version 9.0: high-level integration of gene regulation, coexpression, motif clustering and beyond. Nucleic Acids Research, 44(D1), D133–D143.

Gustavsen et al. (2019). Rcy3: Network biology using cytoscape from within r. F1000Research.

Hahne, F. and Ivanek, R. (2016). Statistical Genomics: Methods and Protocols, chapter Visualizing Genomic Data Using Gviz and Bioconductor, pages 335–351. Springer New York, New York, NY.

Huber, W. et al. (2015). Orchestrating high-throughput genomic analysis with Bioconductor. Nature Methods, 12(2), 115–121.

Lawrence, M. et al. (2013). Software for computing and annotating genomic ranges. PLoS Computational Biology, 9.

Ledezma-Tejeida, D. et al. (2017). Genome-wide mapping of transcriptional regulation and metabolism describes information-processing units in escherichia coli. Frontiers in Microbiology, 8.

Ledezma-Tejeida, D. et al. (2019). Limits to a classic paradigm: most transcription factors in e. coli regulate genes involved in multiple biological processes. Nucleic Acids Research, 47(13), 6656–6667.

Santos-Zavaleta, A. et al. (2018). RegulonDB v 10.5: tackling challenges to unify classic and high throughput knowledge of gene regulation in E. coli K-12. Nucleic Acids Research, 47(D1), D212–D220.

Seshasayee, A. S. et al. (2006). Transcriptional regulatory networks in bacteria: from input signals to output responses. Current Opinion in Microbiology, 9(5), 511–519.

